# Computational algorithms and neural circuitry for compressed sensing in the mammalian main olfactory bulb

**DOI:** 10.1101/339689

**Authors:** Daniel Kepple, Brittany N. Cazakoff, Heike S. Demmer, Dennis Eckmeier, Stephen D. Shea, Alexei A. Koulakov

**Author notes:** **Corresponding author**: Alexei A. Koulakov.

## Abstract

A major challenge for many sensory systems is the representation of stimuli that vary along many dimensions. This problem is particularly acute for chemosensory systems because they require sensitivity to a large number of molecular features. Here we use a combination of computational modeling and *in vivo* electrophysiological data to propose a solution for this problem in the circuitry of the mammalian main olfactory bulb. We model the input to the olfactory bulb as an array of chemical features that, due to the vast size of chemical feature space, is sparsely occupied. We propose that this sparseness enables compression of the chemical feature array by broadly-tuned odorant receptors. Reconstruction of stimuli is then achieved by a supernumerary network of inhibitory granule cells. The main olfactory bulb may therefore implement a compressed sensing algorithm that presents several advantages. First, we demonstrate that a model of synaptic interactions between the granule cells and the mitral cells that constitute the output of the olfactory bulb, can store a highly efficient representation of odors by competitively selecting a sparse basis set of “expert” granule cells. Second, we further show that this model network can simultaneously learn separable representations of each component of an odor mixture without exposure to those components in isolation. Third, our model is capable of independent and odor-specific adaptation, which could be used by the olfactory system to perform background subtraction or sensitively compare a sample odor with an internal expectation. This model makes specific predictions about the dynamics of granule cell activity during learning. Using *in vivo* electrophysiological recordings, we corroborate these predictions in an experimental paradigm that stimulates memorization of odorants.

## INTRODUCTION

In order to survive, organisms navigating the sensory world need to extract as much information as possible from their environment. The bandwidth of initial sensory processing by peripheral receptors constitutes a critical bottleneck for the nervous system’s ability to extract and represent high-dimensional information from the environment (Barlow, 2001; Lorincz et al., 2012). Chemosensation is a particularly acute example. The chemicals an organism can potentially encounter in its lifetime vary along many dimensions, and they are also vast in number, thus presenting an enormous challenge. How does the brain maintain specificity and accuracy in detection of environmental chemicals, while also representing as many chemicals as possible?

One solution to this problem is to have an array of highly-specific molecular detectors that uniquely signal the presence of one or a small number of highly important compounds (Ai et al., 2010; Kurtovic et al., 2007; Semmelhack and Wang, 2009; Suh et al., 2007; Suh et al., 2004). This ‘labeled line’ scheme allows completely independent control over the response to individual odors, however the number of chemicals that can be represented is limited to approximately the number of receptor types. The mammalian main olfactory system appears to have evolved a different solution. That system uses a large number of receptor types that are relatively broadly tuned to the presence of chemical structural elements or molecular features (Araneda et al., 2000; Buck and Axel, 1991). Consequently, each receptor responds to multiple chemicals, and each chemical activates multiple receptors (Malnic et al., 1999). In this case, the representational bandwidth for odors is much larger, but odor-specific learning is complicated by the interdependence of the representations of odors that activate overlapping sets of receptors.

We propose that the mammalian main olfactory bulb (MOB) solves this problem by implementing a ‘compressed sensing’ algorithm. Compressed sensing exploits the sparseness of input signals to successfully recover them from lower bandwidth representations (Baraniuk, 2007; Donoho and Tanner, 2005, 2006). Relative to the large number of potential chemical features, the specific set of features that constitute a given odorant is sparse, presenting an opportunity for the olfactory system to take advantage of compressed sensing. In practice, we show that this enables the MOB to efficiently represent and recognize on the order of 10^7^ distinct monomolecular compound with only ~1000 receptors.

Koulakov and Rinberg (2011) have developed a Sparse Incomplete Representations (SIR) model of the olfactory bulb, wherein mitral cells, which carry information on to deeper brain targets, effectively compute the difference between active sensory inputs and a learned template encoded in the inhibitory granule cells. Consequently, mitral cells are predicted to signal the difference between the actual stimulus and an internal expectation, as opposed to directly and explicitly signaling the stimulus itself. We propose that learning shapes inhibitory inputs into the mitral cell to achieve the ideal balance with their receptor-dependent inputs. Thus, granule cell representations approach negative ‘mirror images’ of the corresponding receptor activation.

We extend the SIR model to show how the mitral-to-granule cell network can implement a compressed sensing algorithm. The combinatorial pattern of activation of olfactory receptors carries a compressed representation of sparse vectors of molecular concentrations. By virtue of their vastly greater number, and their sparse connectivity, granule cells are able to decompress the pattern of receptor activation. In other words, granule cell activities are predicted to accurately recover information about concentrations of individual independent mixture components. Any mismatch of the decompression result is represented in the residual activities of mitral cells. The olfactory bulb circuit is uniquely suited to implement this algorithm, however the dendrodendritic mitral-to-granule cell connections must have the capacity to learn the receptor affinities to the individual mixture components.

Here, we derive, investigate, and experimentally test the predictions of a learning rule for storing information about olfactory receptor binding affinities in the strengths of dendrodendritic synapses. Incorporating this learning rule into the SIR model confers several extremely useful properties. First, we show that our model stores highly efficient representations of learned odors by competitively selecting a sparse basis set of “expert” granule cells. Second, we show that our model can simultaneously learn separable representations of each component of an odor mixture without exposure to those components in isolation. Finally, we show that these mechanisms allow a sophisticated adaptation strategy in which responses to select groups of mixture components can be modulated independently from other components present.

Our model makes several very specific and counterintuitive predictions with regard to granule cell dynamics during odor learning. To test these predictions, we used *in vivo* electrophysiology to observe the dynamics of granule cells during an odor learning paradigm. We previously reported that stimulation of the noradrenergic brainstem nucleus locus coeruleus suppresses mitral cell responses to paired odors and also facilitates learning (Shea et al., 2008). Our data recapitulates the changes in granule cell responses we observe in our model; most cells are suppressed, while a minority show a dramatic increase in firing in response to the learned odor. Despite the widespread decrease in granule cell output, the data also suggest that the mitral cells paradoxically experience a net increase in granule cell inhibition. Our observation of these properties *in vivo* supports the physiological plausibility of key features of our model.

## METHODS

### Animals

We performed experiments on adult (aged 6–12 weeks) male C57Bl/6 mice (Charles River). Mice were maintained on a 12h–12h light-dark cycle (lights on 07:00 h) and received food *ad libitum*. All procedures were conducted in accordance with the National Institutes of Health’s *Guide for the Care and Use of Laboratory Animals* and approved by the Cold Spring Harbor Laboratory Institutional Animal Care and Use Committee.

### Electrophysiology

Adult male sexually naive C57/Black6 mice (Charles River Laboratories) were initially anesthetized with 100 mg/kg ketamine and 5 mg/kg xylazine. Subsequently, anesthesia was maintained with sevoflurane or isoflurane (~1% in pure O_2_). We measured respiration using a foil strain gauge (Omega Engineering) that was placed on the surface of one side of the animal’s abdomen and connected to a bridge amplifier (Omega Engineering). We performed *in vivo*, loose-patch recordings using borosilicate micropipettes (15–25 MΩ) filled with intracellular solution (125 mM potassium gluconate, 10 mM potassium chloride, 2 mM magnesium chloride and 10 mM HEPES, pH 7.2) containing 1.5% Neurobiotin. After each recording, we labeled cells using positive current injection (+500-700 pA; 0.5 Hz) of Neurobiotin for 15–25 min. All neurons were identified as granule cells by directly visualizing the Neurobiotin stained cell, or by unambiguously observing the tip of the electrode track within the granule cell layer. Thirty extracellularly recorded granule cells from throughout the MOB are included here. We also performed *in vivo*, whole-cell intracellular recordings using borosilicate micropipettes (6–10 MΩ) filled with intracellular solution (130 mM potassium gluconate, 5 mM potassium chloride, 2.5 mM magnesium chloride, 10 mM HEPES, 4 mM Na-ATP, 0.4 mM Na-GTP, 10 mM phosphocreatine, and 0.6 EGTA, adjusted to 285 mOsm and pH 7.2) Neuronal spiking and intracellular potentials were recorded using a BA-03X bridge amplifier (npi Electronic Instruments), low-pass–filtered at 3 kHz and digitized at 10 kHz. Data were digitally acquired using Spike2 software and analyzed offline using Spike2 and Matlab.

### Sensory stimulation

Odor stimuli were delivered as described (Shea et al., 2008). Stimuli were selected from a set of monomolecular odors diluted to 1% v/v in mineral oil. An additional 1:10 flow dilution in our olfactometer resulted in a final concentration at the nose of 0.1% saturated vapor. Odorants were delivered to the nose (2 s delivery, 30 s interstimulus interval) via flow dilution into the oxygen stream (10% into 1.5 L/min O2) using a custom 64-channel olfactometer. In some experiments, sensory receptor neurons were directly optogenetically activated in mice that express Channelrhodopsin-2 in all olfactory sensory neurons (OMP-ChR2; Dhawale et al., 2010). In these cases, optogenetic activation was achieved by illuminating the prep with a 473 nm LED (Luminus Devices) connected to a fiber optic pipe (Edmund) for 1 s.

### LC stimulation

Electrical stimuli were applied through monopolar tungsten microelectrodes (0.5–1 MΩ; Micro-Probe) and consisted of 5 s, 5 Hz trains of 200 μs, 50 μA biphasic pulses delivered to locus coeruleus ipsilateral to the recording electrode and beginning 1 s before stimulus onset. First, we measured baseline responses to sensory stimulation (either a single repeated odor or 1 s of 473 nm light) for 20 - 40 trials, and then we paired the sensory stimulation with a train of electrical pulses to LC. We continued to record responses to the sensory stimulus for another 20 - 40 presentations without electrical activation of LC.

### In vivo measurement of feedback inhibition in mitral cells

Recurrent granule cell inhibition in mitral cells was measured *in vivo* as described (Abraham et al., 2010). A train of 20 brief pulses of positive somatic current injection was calibrated in amplitude and duration to reliably evoke 20 individual action potentials (mean pulse amplitude: 289 ± 130 pA; mean pulse duration: 6.0 ± 2 ms). This train was applied to a mitral cell recorded in the whole-cell intracellular configuration from an anesthetized mouse. Trains were triggered by the respiratory signal according to a manually adjusted threshold in order to maintain a consistent phase relationship between current injection trains and the breathing rhythm. Membrane potential was manually held constant as needed with small adjustments to the holding current (mean range of adjustment: 34 ± 30 pA). At least 10 trials were collected, and we measured the integral amplitude over 1.5 s of the ensuing after hyperpolarization (AHP). This electrophysiological event has previously been established to represent the magnitude of feedback inhibition from granule cells. Then, we presented 20 2 s trials of odor, each paired with a 5 s train of 200 ms, 50 μA, electrical stimulation pulses at 5 Hz that began 1 s before the odor and extended 2 s after each of 20 trials of paired odor. Finally, we once again measured the integral amplitude of the post-stimulus (AHP).

### Data analysis

All data were analyzed with Matlab (Mathworks). The times of maximum inspiration and expiration were extracted from the strain gauge signal, and neuronal firing rate was computed for each breath throughout the entire recording for each cell. Odor-evoked response strength for each trial was defined as the mean firing rate during the breaths in which the cell responded (3.86 ± 1.4 breaths; range = 2 – 7) subtracting the mean firing rate during the preceding 10 breaths. LED-evoked response strength was defined as the mean firing rate during the 1 s light stimulus, subtracting the mean firing rate during the preceding 6 s. Changes in response strength were assessed for each experiment with an unpaired t test comparison of the pre-LC stimulation responses with the post-LC stimulation responses. These results were then subjected to an FDR procedure (Benjamini and Hochberg, 1995) to correct for multiple comparisons. For comparison across cells with different firing rates, we transformed the response strength for each trial into a Z-score. The Z-score was calculated relative to a distribution for that cell of firing rates during an equal number of randomly chosen breaths, subtracting the mean firing rate for the preceding 10 breaths.

To quantify changes in the magnitude of the AHP after LC-odor pairing, we first corrected the membrane voltage signal to remove respiratory-coupled subthreshold activity (Abraham et al., 2010). All traces were median-filtered with a 20 ms kernel to remove action potentials. Segments of respiratory-coupled subthreshold membrane voltage traces were taken from between trials and subtracted from the traces of adjacent AHP trials. The corrected traces were averaged for the pre-LC stimulation trials and the post-LC stimulation trials. Odor responses during the pairing phase were calculated from averaged median-filtered traces from all odor trial that were not corrected for respiratory-coupled activity. The magnitude of averaged AHP traces were measured as the integral of the AHP with respect to baseline for 1.5 seconds from the end of the current injection train. The magnitude of averaged odor response traces were measured as the integral of the signal with respect to baseline during the 2 s odor presentation.

## RESULTS

### Sparse odorant coding

In mammalian olfactory systems, volatile odorant molecules are drawn into the nasal cavity by respiration, where they contact and activate a population of receptor neurons (Buck and Axel, 1991). In mice for example, each of these olfactory receptor neurons (ORNs) expresses one of ~1000 distinct olfactory receptor proteins (Chess et al., 1994). The set of odorant molecule binding affinities for the particular receptor protein an ORN expresses determines how that neuron will respond to odors (Araneda et al., 2000; Malnic et al., 1999; Ressler et al., 1994). ORNs expressing the same receptor protein type converge their outputs at a location in the olfactory bulb called a glomerulus, where they make synaptic contact with an exclusive set of mitral cells (Mombaerts et al., 1996). The exclusivity of this connection creates “channels” of mitral cells that convey the activity levels of individual receptor types to the olfactory cortex.

The activity of all of the olfactory receptors in response to an odor is captured by these channels, and this population activity can be described as a vector 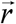 with each element *r_j_* being the activity of the glomerulus with receptor type number *j*. Given that most mammalian systems have ~1000 olfactory receptors, this vector constitutes a complex, high dimensionality representation. Complexity of the neural representation may be useful however, in light of the complexity of olfactory stimulus space. A single molecule can vary along many dimensions, such as the number of atoms, the type of atoms, and their structural arrangement. The complexity of single molecules is compounded by the fact that naturally encountered odors are typically mixtures of many different individual molecules. Consequently, we will henceforth use the term “odor” to denote a mixture of one or more individual odorant molecules.

In this formulation, any odor can be quantitatively described as a list of all of the individual odorant molecules present in the stimulus and their abundance. We therefore define another vector 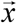, in which each element *x_i_* is the concentration of the *i*-th molecule.

Finally, if we describe the affinity between the molecule *i* and receptor type *j* in a matrix *A_ji_*, we can hypothetically predict the response 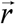 for any arbitrary odor 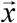. Using the simplest receptor-binding model of a single binding site with no cooperativity, the law of mass action provides a relation between receptor activities *r_j,_* molecular concentrations *x_j_*, and affinities *A_ji_*:

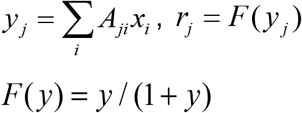

Here *F*(*y*) is the nonlinear function that describes the activation of a receptor. Because the response *r_j_* is related to its input *y_j_* via a simple monotonic function *F*, we can assume that networks analyzing responses of receptor neurons have the linear component of the response *y_j_* available to them. This enables us to think of ORNs as encoding 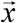 in a vector 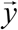 with elements *y_j_* linearly dependent on 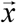.

The length of 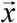 corresponds to the number of available volatile molecules from which a target odor may be composed. This quantity greatly exceeds the number of receptor types in any olfactory system. Therefore, the encoding performed by ORNs, i.e. mapping of the vector 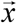 onto vector 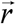, would seem at first to require discarding much of the information content of the odor. Relief comes from the fact that for real world odors, 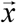 is sparsely occupied. Most odorant molecules are not present in any particular environment, greatly reducing the information content of 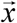. In fact, we can ask how many receptor types an olfactory system would need if its receptor encoding were to fully capture the information of a sparse vector of molecular concentrations 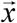. To be able to recover 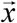, the information content of ORN responses 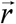 has to match the information contained in 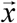. To find the amount of information contained in any vector *H* we can use a formula known from statistical physics

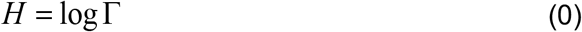

Here Γ is the number of combinations that the values of a given variable can take (here and below, log refers to a binary logarithm). For example, a binary string of length *N* can take Γ = 2^*N*^ values, leading to the information content *H* = log2^*N*^ = *N*, i.e. *N* bits.

If the total number of odorant molecule types is *M*, and the number of molecular components that are typically present in a mixture is *K*, the information content of sparse vector 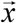 can be estimated from the number of combinations that the vector can take. This quantity 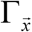 is, for the most part, determined by the identities of nonzero components of 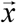, i.e. 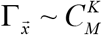,

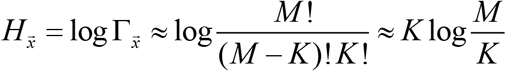

Likewise, we can compute the information capacity of an olfactory system containing *N* receptor types with binary responses as: 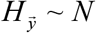. For this type of receptor encoding to capture all of the information in 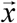, we find the following constraint on the number of receptor types *N*:

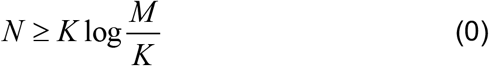

Because *M* is only present in the log, we can accept a wide range of uncertainly in our estimate for the number of odorant molecules.

From looking at the number of molecules under 300 Dalton in PubChem, we estimate the number of odorant molecules to be on the order of 10 million. Psychophysical studies suggest that human observers can detect roughly 12 monomolecular components from a mixture, and therefore we use K~10. This makes our estimate for the number of olfactory receptors necessary to discover components in the mixture *N* ≥ 200. Remarkably, this estimate indicates that the receptor ensembles of both human (*N* ~ 350) and mouse (*N* ~ 1000) have sufficient information capacity to recover the full concentration vector 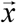.

### Reconstructing sparse odorant stimuli

Since the information capacity of the olfactory stimulus representation 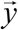 likely well exceeds the information capacity of the stimulus 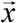, it is possible for the olfactory system to fully recover 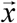 from receptor responses 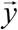. In other words, the brain has sufficient capacity available to recover the complete concentration vector and therefore, the odor identity. A potential algorithm for achieving this comes from the concept of compressed sensing (Baraniuk, 2007; Donoho and Tanner, 2005, 2006). In principle, recovering the million-dimensional vector 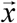 from a thousand-dimensional vector 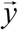, such that 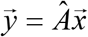, would necessitate solving a system of very few equations with a large number of unknowns. Compressed sensing suggests that this can be accomplished if 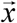 is sparse by finding the solution with the minimum *ℓ*_1_ norm, i.e. the smallest, most parsimonious solution. Thus, the olfactory system is expected to solve the following problem:

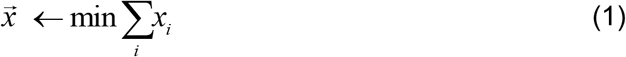

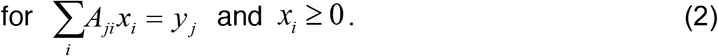

The latter inequality imposes the requirement for a solution with only non-negative elements, because concentrations of individual molecules cannot be negative. How can the brain solve this problem? Instead of solving this problem exactly, we can consider solving an equivalent ridge regression problem

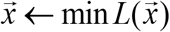

where

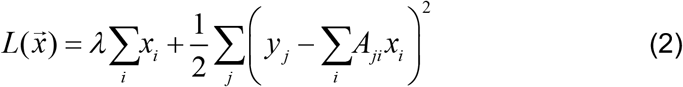

The minimum is expected to be found for the vectors the *ℓ*_1_ norm of 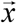 with non-negative components, i.e. *x_i_* ≥ 0. The first term in the function 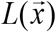 here describes the solution of the equation 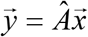, while the second term is the error of the solution. When 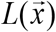 is minimized a compromise is found between these two constraints, i.e. vector 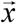 that satisfies the equation 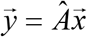 reasonably well and has a small measure *ℓ*_1_ due to the first term. The coefficient *λ* here describes the relative importance of these two constraints. A small *λ* would make the solution of the equation 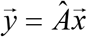 more precise, because the error in solving this equation [second term in equation (2)] is more costly. Thus, when *λ* → 0, the solution of problem (2) will approach the compressed sensing solution (1). Thus, for sufficiently small *λ*, one can substitute solving the compressed sensing problem (2) with the ridge regression problem (2).

How could the olfactory system solve the ridge regression problem (2)? We previously showed (Koulakov and Rinberg, 2011) that the reciprocal circuitry between granule cells and mitral cells in the olfactory bulb allows granule cells to dynamically minimize the following function, also known as the Lyapunov function:

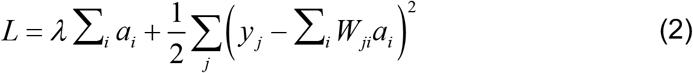

In equation (6), *a_i_* is the activity of granule cell *i, W_ji_* is the synaptic weight between granule cell *i* and mitral cell *j*, and *y_j_* is the receptor input into *j*. In this equation, *λ* is the granule cell firing threshold assumed to be the same for all cells. We also disregarded the term 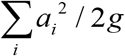, which can be made small by increasing the granule cell activation gain *g*. Note the similarity between equation (2) and equation (2). In comparing the two equations, it is clear that granule cell activity *a_i_* plays the same role in equation (2) as the concentration vector *x_i_* does in equation (2).

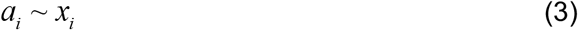

Indeed, in the first term of both equations (2) and (2), both *x_i_* and *a_i_* are under a parsimony constraint. In the second term of (2) and (2), both 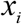 and 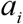 are multiplied by a matrix to minimize the squared error of approximation of *y_j_*. In addition to playing a similar mathematical role, 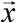 and 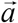 also share a positivity constraint. 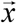, the vector of odorant molecule concentrations, must be positive as negative concentrations are non-physical. Granule cell activities *a_i_*, defined by firing rate, also cannot go below zero. The lengths of the two vectors are similar. The length of vector 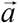 is equal to the number of granule cells, which have been estimated to be on the order of a few million. Our estimate on the number volatile molecules on PubChem, and therefore the length of 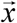, is on the same order. It is therefore plausible that the mitral-granule cell network in the olfactory bulb can perform the reconstruction (2), with the ensemble of granule cell activities representing the brain’s reconstruction of 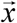, the concentration vector of the odor stimulus.

### The granule cell network can reconstruct odor stimuli

The structure and dynamics of the granule-mitral cell network suggest the possibility that this network performs sparse reconstruction of the concentration vector 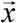. To make this possible, the synapses between granule and mitral cells, *W_ji_*, should learn the affinity matrix *A_ji_*.

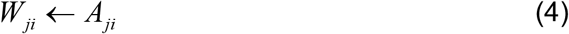

In that case, the granule cell network Lyapunov function (2) and the ridge regression reconstructing the concentration vector (2) are identical. Therefore, in this paper we investigate how the synapses between granule and mitral cells, *W_ji_*, could learn the affinity matrix *A_ji_*.

In our previous work (Koulakov and Rinberg, 2011), we showed that granule cell activities minimize the Lyapunov function given by equation (2), thus following a form of gradient descent of the cost function. To find the learning rules for network weights *W_ji_* we can use a similar principle. To find the learning rule for synapses *W_ji_*, we use stochastic gradient descent on the Lyapunov function with respect to the synapse weights *W_ji_*, yielding:

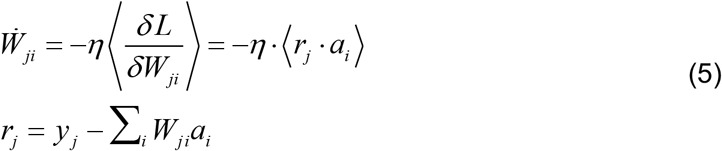

Here, *η* is the learning rate and *r_j_* is the activity of the mitral cell *j*, and 〈…〉 denotes averaging over an ensemble of stimuli. Because the update rule depends only upon the activities of mitral cell *j* and granule cell *i*, it can be performed locally within each mitral cell and granule cell synapse in a Hebbian fashion. According to this learning rule, synapses of active granule cells will change until 〈*r_j_* · *a_i_* = 0, which is possible if *r_j_* = 0 for every odor, or 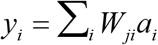.

If the ensemble of granule cell activities represents the reconstruction of the concentration vector 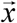, then this can be rewritten as 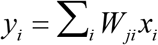. Because, by definition, 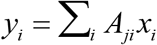, it follows that granule cells will change until 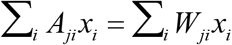 and therefore *W_ji_* = *A_ji_*. That means that a biologically plausible local learning rule between granule and mitral cells would produce a network that reconstructs a representation of the odor stimulus by the granule cells.

For the reverse synapse, 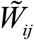, from mitral cells to granule cells, we use a similar learning rule

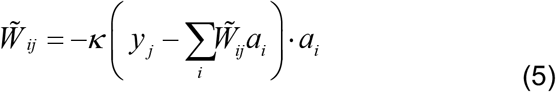

These two learning rules, (5) and (5), guarantee that the forward and reverse synapses are proportional, which is sufficient for the Lyapunov function (2) to apply.

### The learning rule for corticobulbar synapses

We also propose learning rules for corticobulbar synapses onto granule cells. We argue that these synapses define the overall firing threshold for granule cell *i*, *λ_i_* which can vary across cells and is experience-dependent. With this assumption, we can modify net cortical input for each granule cell without explicitly including cortical cells in our network. If we rewrite the cost term of the Lyapunov function (the first term of equation 3) to depend on *λ_i_*. Then the Lyapunov function becomes:

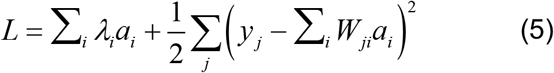

As in equation (5), we find the learning update rule by taking the derivative of the Lyapunov function: 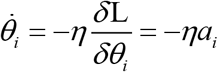. This rule decreases the thresholds of highly active granule cells more rapidly than those of less active cells. As a result, learning is expected to cause these granule cells to increase their odor-driven firing, and to thereby minimize mitral cell responses more efficiently.

### The emergence of expert granule cells

The dynamics of granule-mitral cell networks described by the Lyapunov function (5) and learning rules defined by equation (5) have a simple geometric representation, which we will now describe. Let us assume that the thresholds for granule cells firing are very small, due for example to the sort of learning described above. Minimizing the Lyapunov function (5) with respect to the set of granule cell activities *a_i_*, forces the second term, 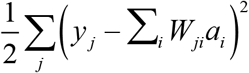, towards zero. This term is an *ℓ*_2_ norm of mitral cell outputs, or the difference between their excitatory inputs, 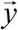, and inhibitory input from granule cells 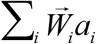. The dynamics of the mitral-granule cell network, therefore, force mitral cell responses towards zero. This, consequently, encourages the inhibitory inputs from granule cells 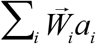 to match the excitatory stimulus input onto mitral cell 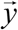, thus establishing a balance between excitation and inhibition by the inputs of mitral cells.

The contribution of granule cells can be visualized using a set of vectors 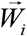. Each of these vectors corresponds to a particular granule cell number *i* and is defined by all the weights of its synaptic contacts with mitral cells. The number of elements in vector 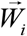 is therefore equal to the number of mitral cells, while the total number of these vectors is equal to the number of granule cells. The ensemble of weight vectors 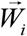 is simply another way to represent the mitral-to-granule cell connection weight matrix 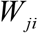. Because the number of elements in each vector 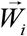 is equal to the number of mitral cells, we can visualize the ensemble of 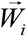 vectors in the same space as the mitral cell input 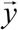 (Figure 1A). The representation of the granule cell input onto mitral cells is a linear combination of these 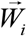 vectors, 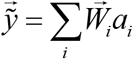 (Figure 1B). From equation (5), as described above, the dynamics of the granule cell network drive this vector to match 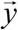.

The requirement for granule cell activity to be positive constrains the set of possible solutions for reproducing 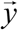. In other words, since *a_i_* ≥ 0, the granule cell reconstruction can only be formed from positive combinations of 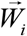. Geometrically, this means that the reconstruction 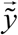 is constrained to the cone with edges 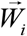 (shaded region in Figure 2B). For inputs 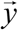 inside the cone, granule cells can build an exact representation of vector 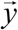, i.e. 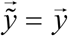, barring corrections introduced by costs in equation (5). In this case, the activity of mitral cells is zero, i.e 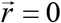 [equation (5)]. If the input into mitral cells from receptor neurons 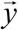 is outside the cone, the representation of 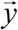 formed by the granule cells is given by the vector nearest to 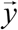 on the cone’s hull. Only the vectors forming the corresponding face of the hull (blue in Figure 2B) are needed to represent this best possible solution. In this case, the unmatched mitral cell output 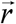 r represents the error of the solution, or the distance between the ideal solution 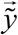 and the closest conic hull vector 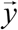. Only granule cells corresponding to vectors forming the nearest hull face will contribute activity to the representation. As a result, population responses of granule cells will be sparse. Because the representation of inputs 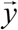 by the granule cells is frequently incomplete, i.e. 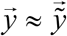 and 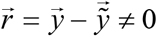, we called this model the Sparse Incomplete Representations or SIR model (Koulakov and Rinberg, 2011).

**Figure 1.**
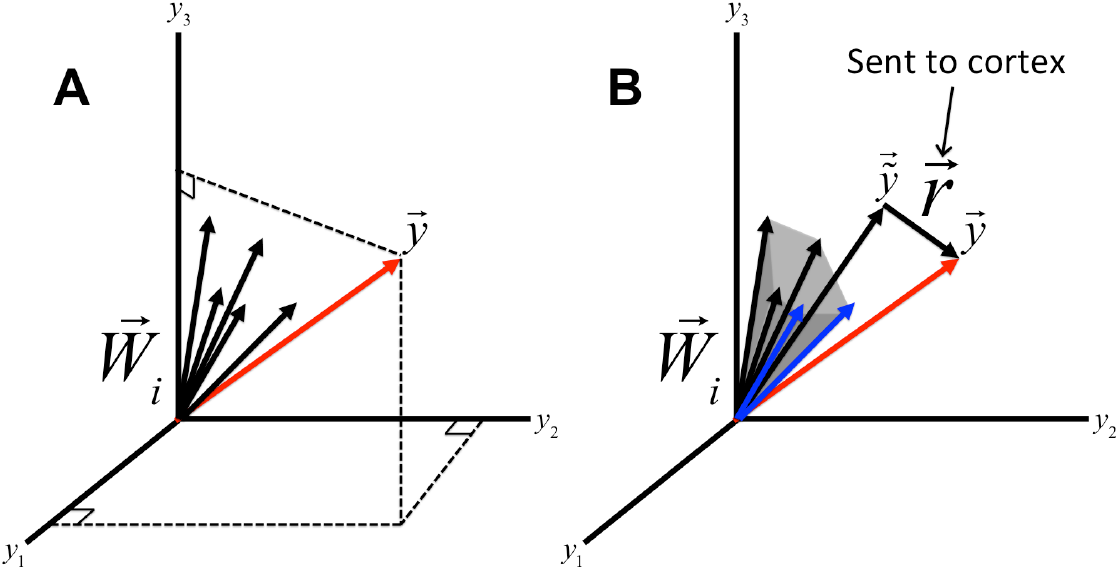
Sparse incomplete representations. This example describes an olfactory system with three mitral cells and five granule cells. (A) Vector 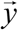 (red) represents the inputs received by mitral cells from receptor neurons. Components of vector 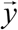 represent inputs from individual glomeruli. The set of vectors 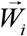 (black) represents synaptic strengths of individual granule cells, numbered by index *i*, with all mitral cells. (B) The granule cell representation of mitral cell input, 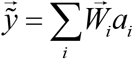, is limited to the convex cone formed by 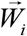 vectors (grey shaded area), due to non-negativity of granule cell activities *a_i_*. For vectors 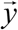 outside the cone, only the blue vectors 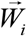, which constitute the nearest face of the cone, are active in the representation. The difference between the true mitral cell inputs 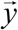 and the granule cell representation 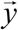 is contained in the vector of mitral cell responses 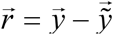 and is sent to the cortex.

**Figure 2.**
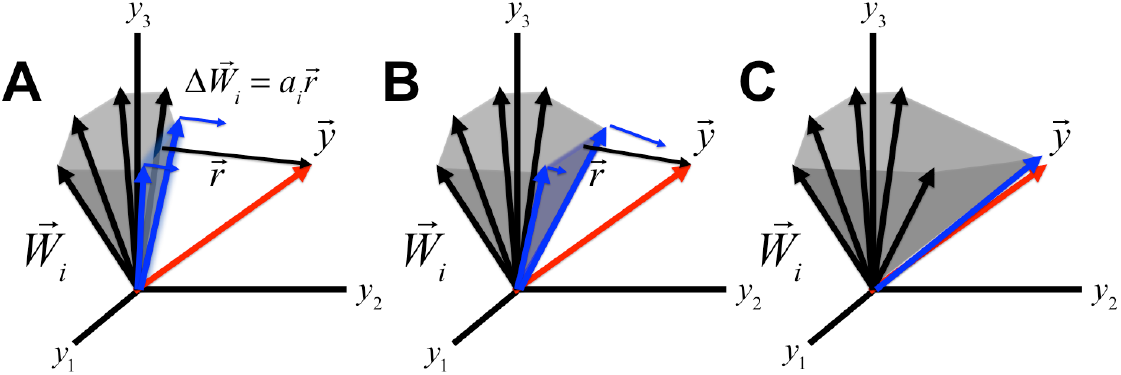
Learning in granule cell synapses. (A) Synaptic weight vector of granule cell *i*, 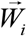, will only change if the cell responds to the given odorant, i.e. is on the face of the cone closest to the mitral cell input 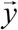. The direction and magnitude of the weight vector update (small blue arrows) are determined by the responses of the mitral cells 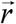 and granule cell activity respectively. (B) As a result, the vector 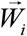 nearest to 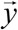 will change the most. (C) After several rounds of updates, a single “expert” granule cell dominates the representation of 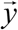.

This geometric formulation of our model can incorporate weight updates specified by the learning rule (4) as well. The learning rule (4) postulates that vectors 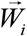 themselves can change. The direction of change is determined by the vector of mitral cell responses 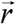. The rate of change is proportional to the response magnitude of a given granule cell *a_i_*. Thus, only weights of granule cells with non-zero responses to the odorant will be updated and the weights of more active cells will be updated more drastically. Since responses of granule cells are sparse, only a small population of granule cells is involved in learning each odorant. These granule cells are situated on the face of the cone that is nearest to the mitral cell input vector 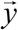 (Figure 2A). Interestingly, the nearest 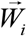 will change the most, as its respective granule cell activity will be strongest. Therefore, with each successive iteration, the vector 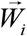 nearest to 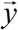 will increasingly dominate the representation (Figure 2B). As this process continues, one granule cell with synaptic weights 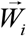 will become the sole contributor to the representation of 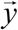 (Figure 2C). We refer to this granule cell as an “expert” granule cell with respect to representing the odor. In practice, this correspondence of one granule cell to one input vector 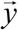 is unlikely because no single granule cell will have synaptic access to all mitral cells. As a result, the degrees of freedom for each 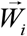 will be limited to the number of mitral cells accessed by granule cell *i*. Therefore, we predict that: 1) The activation pattern evoked by real world odor requires the existence of a minority of strongly responsive “expert” granule cells, and 2) Overall granule cell population activity will be sparse.

### Learning the representations of simple odorants

We next wished to confirm the plausibility of a small cohort of granule cells learning to represent an odorant. Therefore, we devised a model simulating the learning dynamics of the MOB in response to distinguishable patterns of receptor inputs. These patterns were chosen from a commonly used set of visually distinguishable images, six of which are shown in Figure (3A). These were intended to each represent an input pattern associated with a simple odor. The prediction of the model outlined above is that, after learning, each pattern will be represented by a sparse subset of active granule cells that we call “experts”.

**Figure 3.**
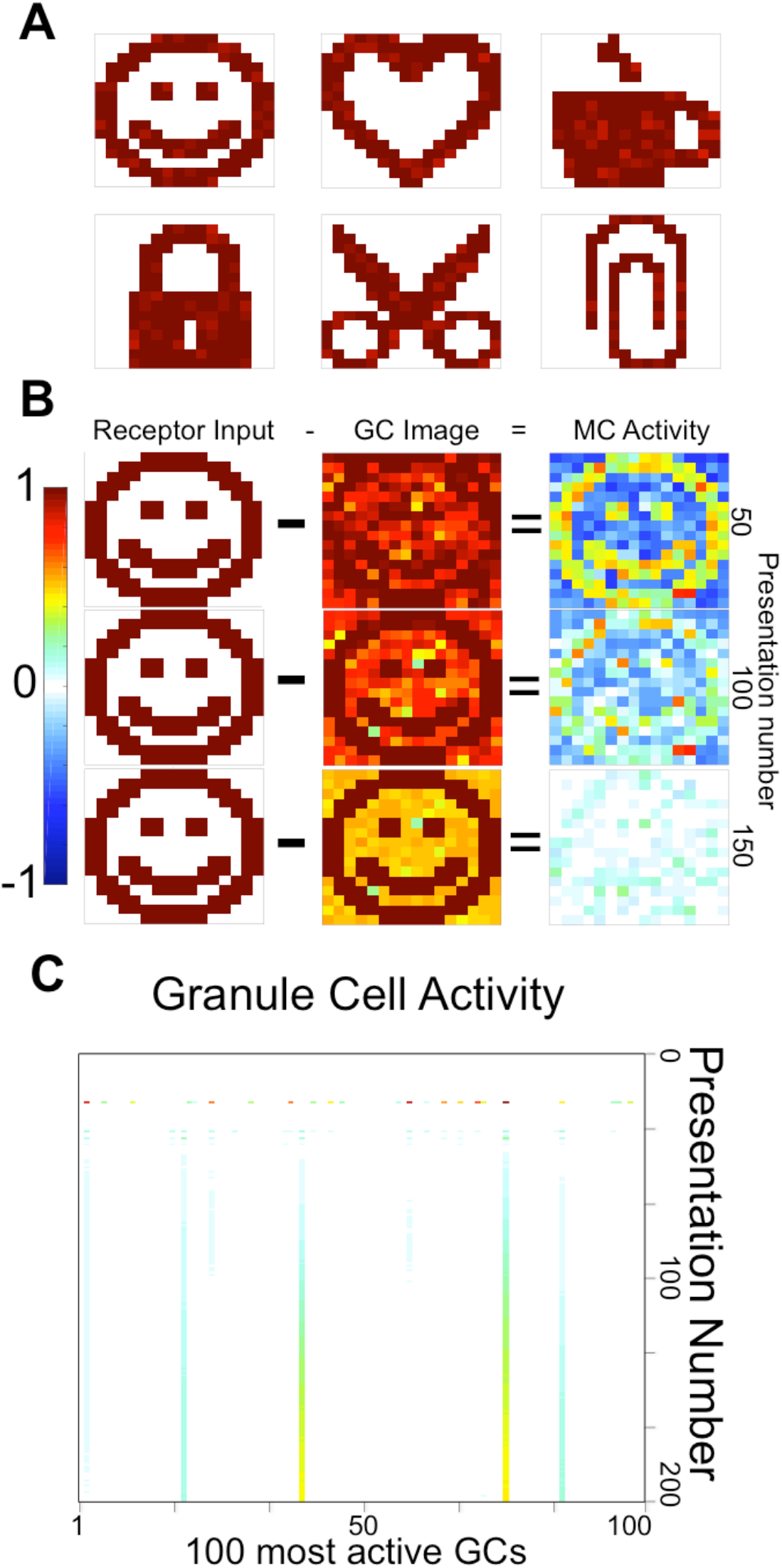
Simulation of granule cell learning. We simulated mitral cell receptor inputs as recognizable images (A). The difference between the receptor input and the granule cell representation is the resulting mitral cell activity. As learning progresses, mitral activity sparsifies and the GC representation becomes more accurate (B). Mean granule cell activity decreases during learning, while expert granule cell activity goes increases.

Our simulations show that, when an odor is first presented, mitral cell outputs closely resemble their inputs (Figure 3B). However, with repeated presentation of a single input 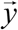, mitral cell responses decrease as the granule cell representation becomes more accurate (Figure 3B). A group of granule cells is initially active, corresponding to the nearest face of the conic hull of GC-MC synaptic weight vectors 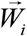 (Figure 2A). In later presentations, most of the granule cell activities drop to zero (Figure 3C). Yet, a small group of granule cells becomes increasingly active. These sparse, highly responsive granule cells closely match our predicted sparse population of “expert” cells. Despite the fact that only a few granule cells are active, the resulting granule cell representation is very close to the mitral cell receptor input, resulting in a weak mitral cell response after learning (Figure 3B). We conclude that granule cell weights can learn the representations of simple odors in mitral cell inputs within an unsupervised learning paradigm.

### Learning components of mixtures

Real world odor scenes are complex and dynamic. Most odors are mixtures in which multiple components are rapidly fluctuating due to turbulent airflow or changing distance to odorant sources. Because the relative proportions of these components are variable, representing odor stimuli with expert granule cells tuned to narrow and specific input configurations presents a challenge. For example, in the extreme case, if each odor stimulus has its own expert granule cell, the network would need one granule cell for each possible mixture. This would be very inefficient and impractical in light of the fact that the number of possible mixtures greatly exceeds the number of granule cells. It would be much more efficient for expert granule cells to learn representations of individual mixture components. In that case, the network would represent exponentially more stimuli using linear combinations of granule cells. Therefore, we extended our simulations to test whether the learning rules described above can enable the network to learn components of mixtures, despite having never been exposed to them in isolation.

We trained the network with complex odors composed of linear combinations of components drawn from an ensemble of input patterns similar to the images shown in Figure 3A. Each pattern was intended to represent an individual simple odorant that can potentially be present in the environment. Out of the nine components, in each presentation, we selected two (Figure 4B) or three (Figure 4C) images to form a mixture. Each component was added with a random coefficient to simulate a fluctuating odor environment. Thus, at no time, was an individual simple odor component presented to the network. Moreover, randomly generated mixtures were never repeated. Then we measured the accuracy of the representations of the individual components.

**Figure 4.**
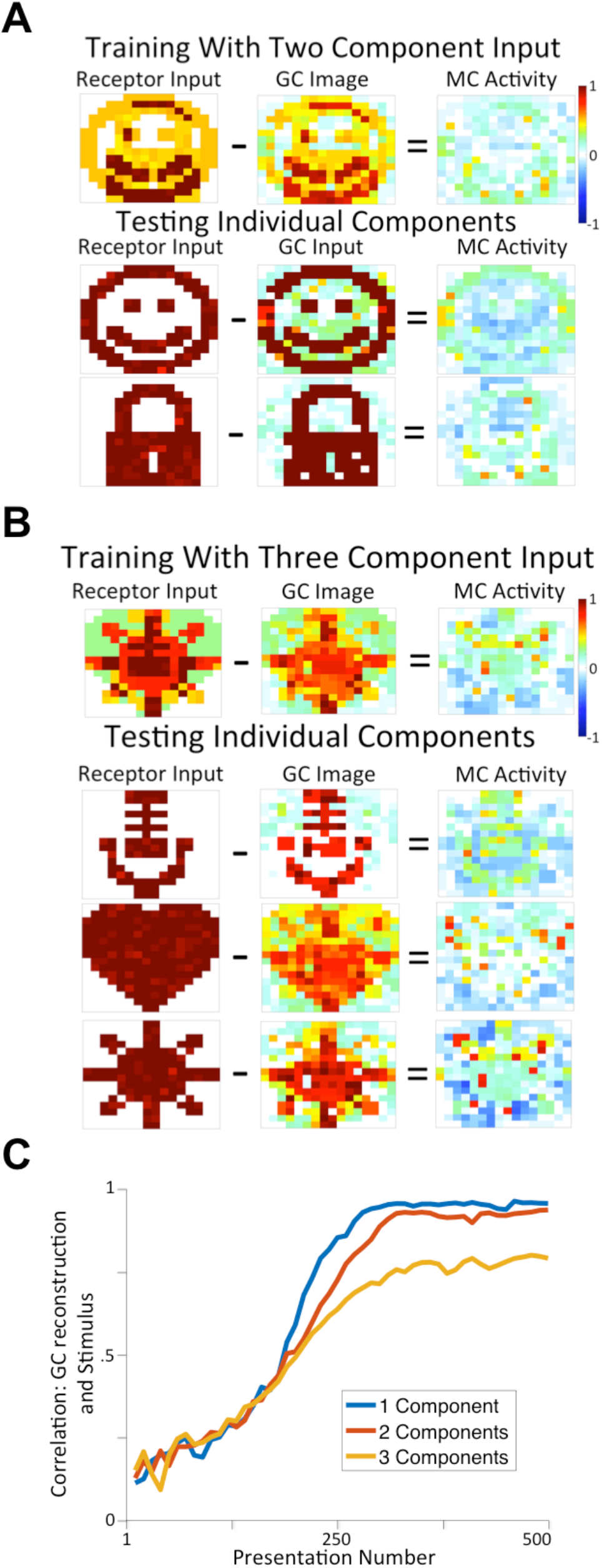
The model can learn components from mixtures. Training on inputs generated from adding, with random coefficients, two randomly selected images from nine component images. After 500 iterations of training, we tested the model on the component images alone (A). We used the same procedure for inputs with three components (B). After each iteration, we presented the nine component images individually, calculating the mean correlation between the input and GC representation across all nine images. Increasing the number of components decreases the mean correlation (C).

We found that the network learns to represent the complex odorants despite the fact that each of the mixtures presented to the network is unique. The network is able to successfully represent mixtures because inhibition from granule cells, at the end of training, almost precisely cancels mitral cell inputs, leading to weak responses of mitral cells to odorants. We tested this for both two- and three-odorant mixtures (Figures 4A and B respectively). We find that, after several presentations, the network learns to accurately reproduce the nine component images despite having never encountered them alone. We conclude that the learning rules described above enable the model network to represent mixtures as linear combinations of learned representations of their individual components. This shows that our model is not just mirroring odorant inputs but is instead able to extract and learn the components in a dynamic stimulus.

### Granule cell dynamics during odor learning

These simulations make two surprising predictions about changes in the activity of granule cells during learning. First, our model predicts that learning sparsens the population of granule cells that respond to the learned odor. This results from enhancement of the output of a minority of ‘expert’ granule cells, and the suppression of the output of other granule cells. Second, our model predicts that despite a widespread reduction in many granule cells’ output, there is an increase in net inhibitory input to mitral cells that respond to the learned odor. We used *in vivo* extracellular and intracellular electrophysiology methods in granule cells and mitral cells to test these two predictions. During our recordings, we facilitated synaptic plasticity and learning associated with odors by electrically stimulating a noradrenergic input to the olfactory bulb.

In two previous studies, we showed that stimulating the noradrenergic brainstem nucleus locus coeruleus (LC) while simultaneously presenting an odor, triggered long-term changes to the responses to that odor (Eckmeier and Shea, 2014; Shea et al., 2008). For example, pairing LC stimulation with an odor in anesthetized mice causes a consistent and enduring decrease in the responses of mitral cells. This firing decrease is dependent on circuitry and noradrenaline release in the olfactory bulb. Importantly, although the mice were anesthetized, in subsequent behavior experiments, mice that are presented with the odor that was paired to LC stimulation exhibit signs that they remember that odor (Eckmeier and Shea, 2014; Shea et al., 2008). Therefore, LC-mediated noradrenaline (NA) release facilitates memorization of odors by modifying olfactory bulb circuits. We therefore used this approach to observe the dynamics of granule cell firing rates during odor learning, We first used ‘loose patch’ *in vivo* electrophysiology methods to record individual granule cells in the main olfactory bulb of anesthetized adult male mice (Cazakoff et al., 2014). Responses were evoked in the granule cells either with 2-second pulses of monomolecular odors, or with 473 nm light activation in mice expressing the optogenetic tool Channelrhodopsin-2 (ChR2) in olfactory sensory neurons (Dhawale et al., 2010). We measured responses to odor or light stimulation (see Methods) for 20 - 40 trials, and then we monitored changes in the response as we paired each sensory stimulus with a brief train of electrical pulses to LC for another 20 - 40 trials, and beyond. Data from several example experiments are depicted in Figure 5.

**Figure 5.**
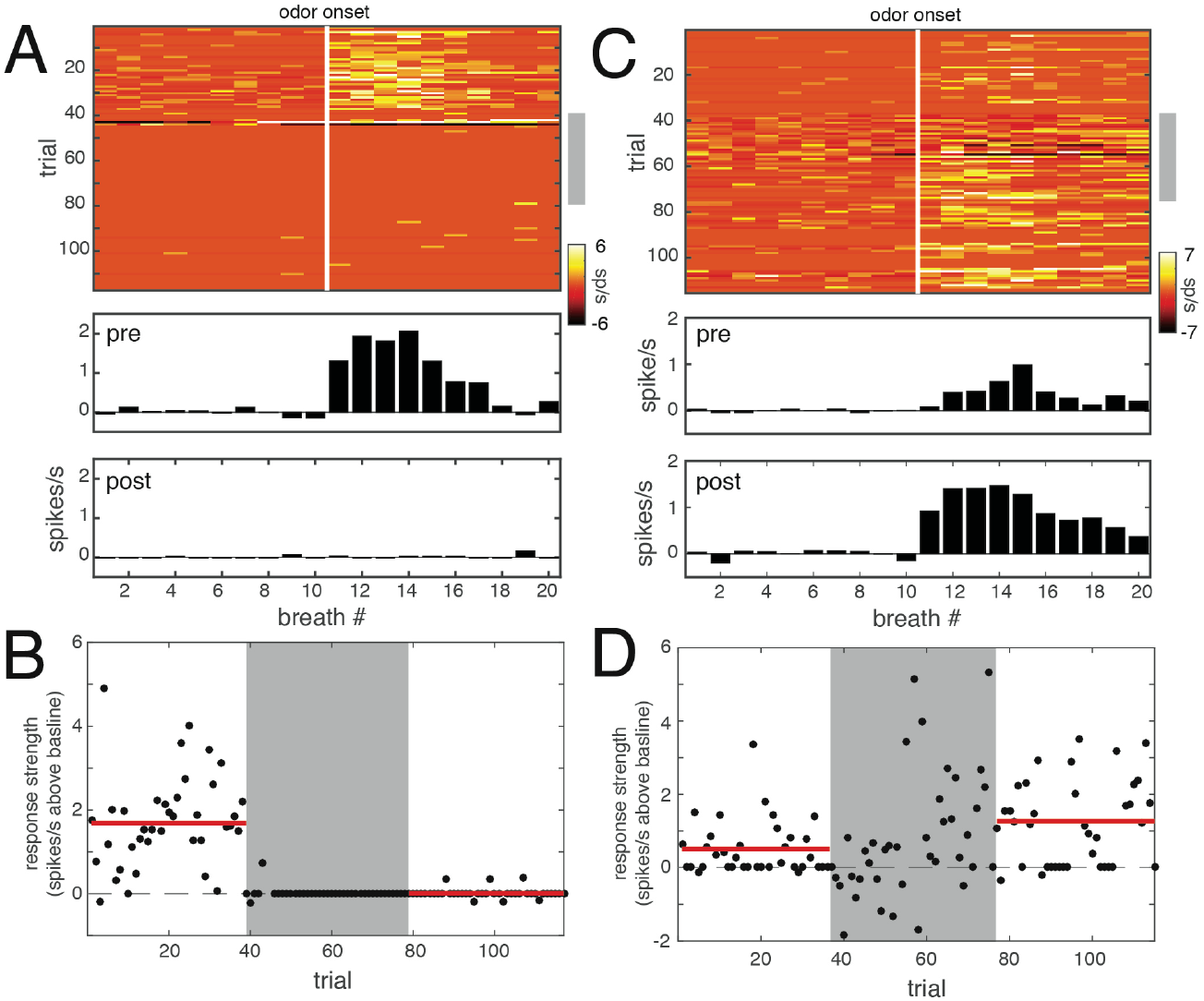
Repeatedly pairing odors with electrical stimulation of the noradrenergic brainstem nucleus locus coeruleus (LC) leads to either long-term suppression or longterm enhancement of sensory responses in granule cells. (A) Two-dimensional peristimulus time histogram (PSTH) depicting the response of an individual granule cell to 117 consecutive 2 s trials of odor presentation. Each row represents one trial, and each bin in the row represents the firing rate (spikes/s) for one respiratory cycle. The vertical white bar denotes the onset of the first breath after the odor is presented. The gray bar on the right indicates that for trials 40-79, each odor presentation coincided with a 5 s, 5 Hz train of 50 μA electrical pulses applied to the ipsilateral LC. The train began 2 s prior to odor onset. The lower panels are PSTHs that depict the mean spike rate response for trials prior to the LC-odor pairing (‘pre’) and for trials after the odor LC-pairing (‘post’). (B) Scatterplot depicting the trial by trial response strength over the whole experiment, computed as the mean firing rate during the stimulus minus the mean baseline firing rate. The gray shaded region denotes the LC-odor pairing trials. The horizontal red bars indicate the mean firing rates for the pre and post pairing epochs. (C, D) Data from a different granule cell that showed increased odor responses after LC-odor pairing over a total of 115 odor trials, organized as in A and B.

Figure 5A and 5C are 2-dimensional peristimulus time histograms (PSTHs) that depict the activity of two different granule cells in response to more than 100 trials of odor presentation. The lower two panels show PSTHs of the mean baseline-subtracted firing rate before and after LC-odor pairing. Most commonly, sensory responses of granule cells were dramatically suppressed by LC-odor pairing, as in Figure 5A. A smaller population showed robustly increased firing in response to sensory stimulation, as in Figure 5C. Figure 5B and 5D are scatter plots of the trial by trial response strength in each experiment, computed as spikes/s above baseline during the response (see Methods.)

As our model predicted, the data from granule cells show that their responses sparsen after pairing sensory stimulation with LC-activation. Sparsening occurs because the majority of cells show suppression of responses, and a minority show strengthened responses. In total, we collected similar data from 20 granule cells, and the results of these experiments are depicted in the scatter plot in Figure 6A. Each point in the plot represents the mean pre-LC pairing sensory response strength compared with the mean post LC-pairing sensory response strength, each expressed as Z scores. In 12 experiments, we collected responses to odor presentation, and in the remaining 8 experiments, responses were collected to optogenetic activation of OSNs. Filled points denote experiments where the post-LC pairing responses were significantly different from the pre-LC pairing responses, as assessed with a two-tailed unpaired t-test (*p* < 0.05), subject to an FDR analysis correcting for multiple comparisons (Benjamini and Hochberg, 1995). Of 20 LC-pairing experiments, 15 showed significant changes in response strength after LC-pairing, including 11 experiments in which responses were suppressed and 4 experiments in which responses were increased. Notably, 3 of the sites that did not change after LC pairing were all 3 sites which responded to the odor with firing suppression, suggesting that LC activity only affects positive, excitatory responses in granule cells. Changes in response strength were uncommon in experiments in which odor responses were measured before and after sham LC stimulation (4 experiments) or before and after LC stimulation was paired with an odor blank (6 experiments) (Figure 6B). Of these 10 cases, 3 showed significant changes in granule cell response strength (1 decrease and 2 increases.) Figure 6C is a histogram of the ratio between post-LC pairing responses and pre-LC pairing responses for 27 experiments in which the pre-LC pairing sensory responses were excitatory, comparing LC pairing sites (red) to control sites (blue.) These data show that noradrenaline release in the olfactory bulb, which facilitates odor memories, results in a bimodal distribution of effects characterized by predominantly dramatic response suppression, but with strong response enhancement at a minority of sites. This result therefore matched the predicted sparsening of granule cell responses during learning.

**Figure 6.**
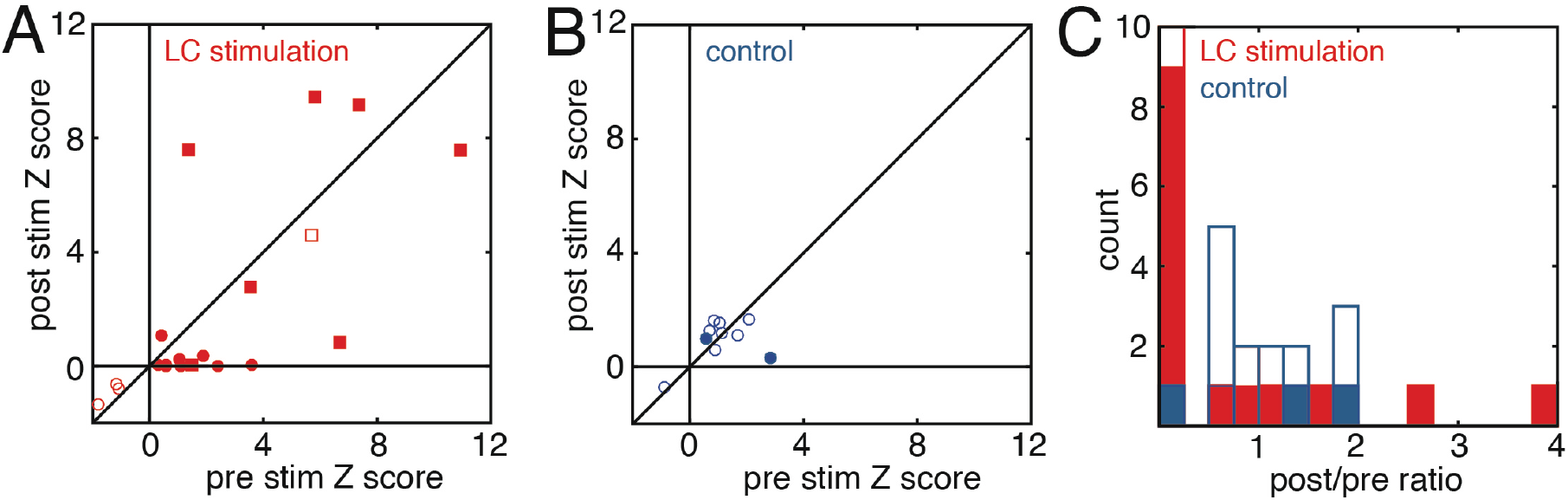
Repeatedly pairing sensory stimuli with electrical activation of LC sparsens the response of granule cells to paired stimuli. (A) Scatterplot of data from 20 experiments in which activation of OSNs by either odors or ChR2 stimulation were paired with LC stimulation. Each point in the plot represents the mean pre-LC pairing sensory response strength compared with the mean post LC-pairing sensory response strength, expressed as a Z score (see Methods). Circles denote responses to odors and squares denote responses to optogenetic activation of OSNs. Filled points designate experiments where the post-LC pairing responses were significantly different from the pre-LC pairing responses, as assessed with a two-tailed unpaired t-test (*p* < 0.05), subject to an FDR analysis correcting for multiple comparisons. (B) Scatterplot of data from 10 experiments in which either sham LC stimulation was performed or LC stimulation was paired to a blank odor. The panel is organized as in A. (C) Histogram of the ratio of postpairing response strengths to pre-pairing response strengths for all experiments in A and B. Red bars denote LC pairing experiments and blue bars denote control experiments. Filled bars designate significant changes, as identified in A and B.

The second prediction of our model is that although most granule cells are suppressed, the net inhibition onto mitral cells that represent an odor is increased after learning. We used whole-cell intracellular recordings from mitral cells to show that pairing an odor with LC stimulation increases inhibitory feedback onto mitral cells that are depolarized by the odor. Previous work has shown that depolarization of mitral cells evokes a sustained ‘after hyperpolarization’ (AHP) that reflects inhibitory synaptic feedback from granule cells (Isaacson and Strowbridge, 1998). Measurement of this event has been used *in vivo* to quantify the strength of granule cell feedback onto mitral cells (Abraham et al., 2010). We therefore used whole-cell intracellular recording methods to determine whether and how LC-odor pairing affects inhibitory synaptic feedback onto mitral cells.

We found that pairing an odor with electrical stimulation of LC led to increased inhibitory feedback from granule cells specifically onto mitral cells that were responsive to the paired odor. Figure 7A depicts the structure of the experiment. The black and red traces depict representative measurements of the integral magnitude of the AHP following a train of 20 action potentials evoked by brief pulses of somatic current injection. The vertical arrowheads are pointing to the AHP in each trace. This measurement was made before and after either 20 trials of an odor paired with electrical stimulation of LC as described above, or an equivalent time during which no electrical stimulation of LC was performed (sham stimulation). Example data in Figure 7B shows that when LC stimulation was coincident with an odor that reliably depolarized the mitral cell, the integral amplitude of the AHP was increased. However, when the mitral cell was not depolarized by the odor, as in Figure 7C, the AHP was unaffected. Figure 7D shows data from a mitral cell that underwent sham stimulation, and the AHP was unchanged. Figure 7E is a scatter plot comparing the net change in the integral area of the AHP following LC odor pairing and sham stimulation. Because the AHP is a hyperpolarizing event, a net negative change in this quantity represents stronger inhibition. The scatter plot in Figure 7F shows that among the mitral cells that were subjected to LC odor pairing, the magnitude of the increase in inhibitory synaptic feedback was significantly correlated with the magnitude of the depolarizing response to the paired odor. This strongly implies that mitral cells responding to the learned odor are specifically targeted for this synaptic modification by NA.

**Figure 7:**
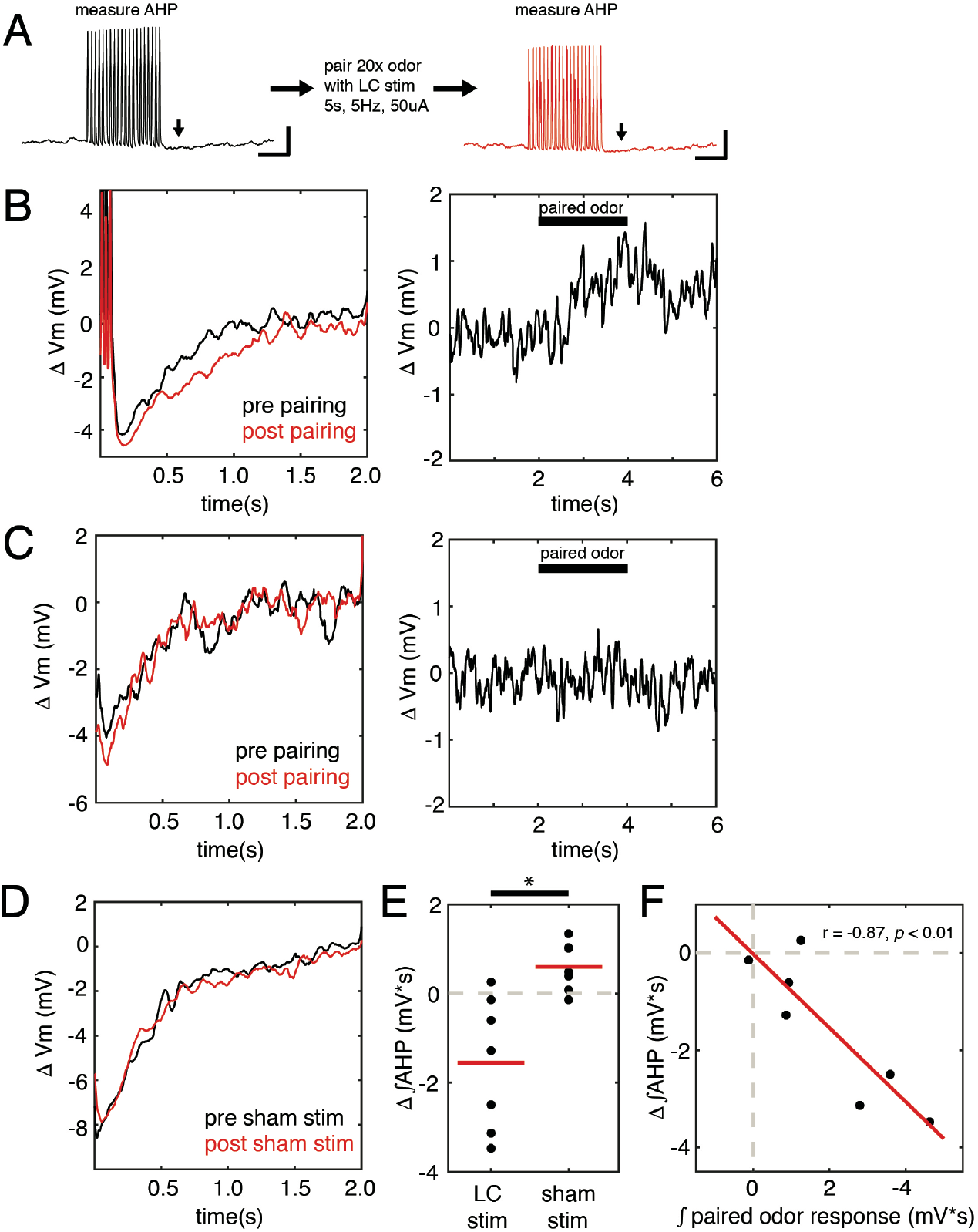
Pairing LC stimulation with odor-evoked depolarization in mitral cells strengthens granule cell inhibitory feedback. (A) Schematic of the experiment. Initially, a train of 20 brief pulses of positive somatic current injection, calibrated in amplitude and duration to reliably evoke 20 individual action potentials, was applied to a mitral cell recorded in the whole cell intracellular configuration from an anesthetized mouse. At least 10 trials were collected, and we measured the integral amplitude of the ensuing after hyperpolarization (AHP). This electrophysiological event has previously been established to represent the magnitude of feedback inhibition from granule cells. Then, we presented 20 2 s trials of odor, each paired with a 5 s train of 200 ms, 50 μA, electrical stimulation pulses at 5 Hz that began 1 s before the odor and extended 2 s after each of 20 trials of paired odor. Finally, we once again measured the integral amplitude of the post-stimulus. (B) Example data from a cell that responded with depolarization to the paired odor and also shows an increase in the integral AHP following odor-LC pairing. Left, the mean AHP prior to odor-LC paring (black) is compared with the same feature after pairing stimulation of LC with a depolarizing odor. Right, the mean subthreshold membrane voltage response to the paired odor. (C) Example data from a different cell that did not respond to the paired odor and also showed no change in AHP. Organized as in (B). (D) Example data showing the mean AHP evoked before and after sham stimulation in LC. The AHP amplitude was unchanged. (E) Scatterplot showing the change in the integral amplitude of the AHP in 14 experiments in which either LC stimulation was paired with an odor or was not performed. The difference between the LC stimulation and sham stimulation groups is significant (unpaired t test, p < 0.01). (F) The magnitude of the change in the AHP amplitude after LC-odor pairing is significantly correlated with the integral amplitude the mean membrane potential response (Pearson’s correlation, r = −0.87, p < 0.01).

## DISCUSSION

Here we have used complementary modeling and *in vivo* electrophysiology approaches to argue that the mammalian main olfactory bulb essentially implements a compressed sensing algorithm. We previously proposed a model for the olfactory bulb in which odor-evoked input patterns in mitral cells are spatially and temporally sparsened by counteracting incomplete suppression from a negative, mirror image representation among the inhibitory granule cells. We termed this representation a ‘Sparse Incomplete Representation’ (SIR) (Koulakov and Rinberg, 2011). Here we extend this model by adding a learning rule that allow the network to adjust its synaptic connections to better represent new odors. This computation allows the bulb to efficiently represent sparse but high-dimensional stimuli. It also makes it possible for the network to learn separable representations of the independently fluctuating components in a complex mixture, without ever encountering them in isolation.

### Dynamics of granule cells during learning

When we observed the dynamics of the granule cells during learning, we observed two paradoxical and seemingly contradictory features. First, as the granule cell representation of an odor stimulus is learned, a large fraction of granule cells drop out of participation in the population response. This emerges in our model as a competitive selection process focuses the network on to an efficient representation by the granule cells that, due to their specific connections, most closely and economically match the stimulus features. We refer to these cells as ‘expert’ granule cells. The broader set of ‘non-expert’ granule cells fall away from this representation. The result of this network dynamic is sparsening of the population response. Notably, when we artificially facilitate odor memorization *in vivo*, a similar sparsening is evident in the data. Specifically, when we pair a sensory stimulus with LC activation, the great majority of granule cells either strengthen their response significantly, or they lose their response almost completely. We note that we observed more apparent ‘expert’ granule cells when we employed broad optogenetic activation, which presumably reflects the increased size of the activated receptor population.

Second, our model further predicts that, despite the loss of granule cell activity, the inhibition sensed by the responsive mitral cells undergoes a net increase. We also observed this pattern in our *in vivo* recordings. We used an established method for quantifying the net inhibitory synaptic feedback onto mitral cells *in vivo* (Abraham et al., 2010). By measuring this quantity before and after LC-facilitated odor learning, we showed that inhibitory feedback was strengthened in mitral cells. Importantly, this was only observed when the test odor evoked a depolarizing response in the recorded mitral cell. Indeed, the magnitude of the change in inhibitory feedback was correlated with the magnitude of the depolarizing odor response. Taken together, these empirical observations are consistent with the granule cell population dynamics we observed in our simulations.

### Computational advantages to compressed sensing of odors

We propose that there are at least three major computational advantages to the neural circuit algorithm we outlined here. First, our model provides an elegant solution for the ‘curse of dimensionality’. Unlike visual stimuli, which are organized topographically in two-dimensional space, and auditory stimuli, which are organized tonotopically according to frequency, odor stimuli occupy a high dimensional space that is defined by their suite of chemical features. While any given odorant will likely possess a relatively small number of these features, the number of potential features these may be drawn from is vast, with each requiring a dedicated sensor. Therefore, representational bandwidth is limited by the number of receptors available. On the other hand, this bandwidth can be expanded with more promiscuous and broadly tuned receptors. The costs of this solution are mitigated by the inherent sparseness of real-world odor scenes, and by the representational capacity of the large network of inhibitory granule cells. Given the sparse set of chemical features present in any odor, granule cells can effectively represent virtually any combination of those features. Thus, the olfactory bulb compresses an efficient representation of sparse odor stimuli that is highly distributed among its array of receptors. The bulb then decompresses the information carried by the receptors by competitive selection of a minimal basis set of granule cells tuned to activity among specific combinations of glomeruli. This feature mitigates the need for an impractically large number of different receptor types.

The second computational advantage to this coding mechanism is that it allows the olfactory bulb to learn the components of an odor without experiencing them in isolation. This capacity is attributable to sensitivity of the granule cells to covariation in concentration among the sets of features that correspond to each component. This is arguably essential for robust odor perception, since real world odor scenes are composed of components that are rarely, if ever, encountered alone.

The third advantage is the capacity for component-specific adaptation or suppression. This network property could be useful in several respects depicted in Figure 8. It could give the piriform cortex (PCx), through its regulation on the granule cell network, control over background subtraction by adapting out the response to irrelevant or expected odors without discrupting the representations of target foreground odors (Figure 8A). Moreover, representing stimuli as mirror images in an inhibitory neural population results in a computation of the difference between the granule cell representation and the sensory input from the glomeruli. Therefore, the output of the olfactory bulb, carried by the mitral cells, signals the error in the granule cell representation. This error signal is highly energetically efficient and allows the olfactory bulb to encode a far greater dynamic range before encountering the biophysical constraints on neuronal firing. Maximizing dynamic range is important in light of the fact that odorants can vary in concentration over many orders of magnitude. Computing a difference error signal is also a highly sensitive mechanism that enables the olfactory bulb to encode very small differences between an expected odor and the actual stimulus.

**Figure 8.**
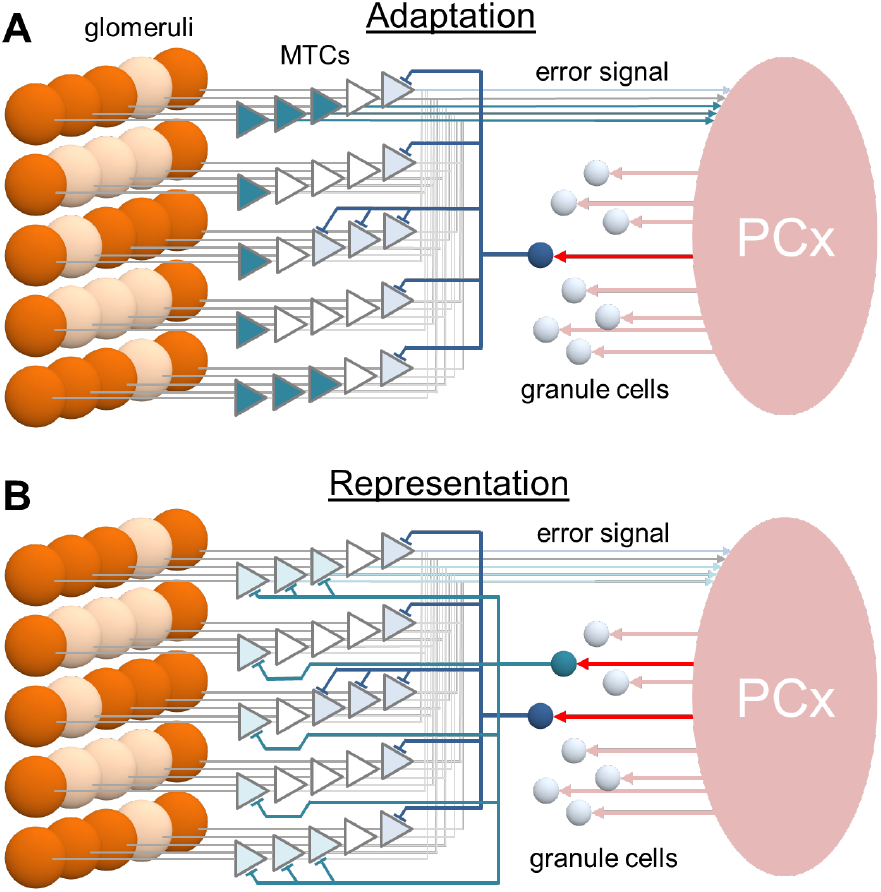
Potential functional significance of sparse odor representations in granule cells for odor recognition. (A) The SIR model may facilitate selective adaptation to known background odorants. In environments where such odorants are known to be potentially present, cortical feedback inputs (red) may lower the threshold for activating the subset of “expert” granule cells responsible for their representation. Thus, the patterns of mitral cell activation by these background odorants are suppressed (light blue). Only the patterns of mitral activation induced by foreground odorants (darker blue) reach the cortex. Thus, the model can remove specific patterns of mitral cell activity associated with odors that are irrelevant for the animal’s behavior. This adaptation mechanism is pattern-specific, leaving mitral cells open for transmitting information about foreground odorants even though their reponses to adapted odors are inactivated by the “expert” granule cells. (B) As an alternative, the SIR model may enable the piriform cortex (PCx) to compare a sample odor against a learned odor expectation. In this mechanism, cortical feedback projections to a cohort of “expert” GCs attempt to match and cancel odorant-induced mitral cell activity. When nearly complete inactivation is achieved, PCx receives weak or negligible inputs, indicates that the odor was identified correctly (no error signal).

### Role of granule cells in odor perception and learning

The results of our modeling and experiments are consistent with the long-standing notion that granule cells figure prominently in odor discrimination and memories. Several lines of evidence support this idea. First, positively and negatively regulating GC activity can respectively improve or degrade performance in a fine odor discrimination task (Abraham et al., 2010; Lepousez and Lledo, 2013; Nunes and Kuner, 2015; Nunez-Parra et al., 2013). Second, odor learning alters rhythmic synchronous population activity (Beshel et al., 2007; Freeman and Schneider, 1982; Martin et al., 2004; Ravel et al., 2003), which appears to critically involve granule cells (David et al., 2015; Kay et al., 2009; Neville and Haberly, 2003; Nusser et al., 2001; Osinski and Kay, 2016; Rall and Shepherd, 1968). Third, noradrenaline-dependent olfactory memories are characterized by stimulus-specific suppression of activity in the mitral cells (Shea et al., 2008; Sullivan et al., 1989; Wilson et al., 1987) and increased GABA release (Brennan et al., 1995; Brennan et al., 1998; Kendrick et al., 1992). For example, NA appears to be important for individual recognition memories that require sensitive discrimination between highly overlapping representations (Brennan and Keverne, 1997). The circuit dynamics we observe in our model and our experimental data suggest that the stimulus-specific suppression of mitral cell odor responses triggered by LC stimulation is achieved through inhibitory feedback from a sparse population of granule cells. Computationally, this implements a subtractive comparison between the odor signature of a given social partner and a memorized template corresponding to a familiar individual. Hypothetically, the result of this computation is the residual mismatch between otherwise highly overlapping representations. Notably, in several studies, the participation of granule cells seems to be of greater importance to this kind of “difficult” discrimination (Abraham et al., 2010; Nunez-Parra et al., 2013).

